# Activation Mechanism and Structural Assembly of the *Mycobacterium tuberculosis* ClpP1P2 Protease and Its Associated ATPases

**DOI:** 10.1101/2025.02.28.640722

**Authors:** Katharina Weinhäupl, Tatos Akopian, Olga Krandor, Dmitry A Semchonok, Rocio Arranz, M.Teresa Bueno-Carrasco, Marcos Gragera, Maelenn Chevreuil, Bertrand Raynal, Yuxin Liu, Jack Lai, WenGen Wu, William Bachovchin, Alfred Goldberg, Eric Rubin, Hugo Fraga

## Abstract

Supramolecular assemblies are integral to cellular biochemical processes, relying on their dynamic nature to fulfill essential functions. The protease ClpP1P2, paired with ATPase partners ClpC1 or ClpX, is vital for the survival of *Mycobacterium tuberculosis* (*Mtb*). While the ClpP1P2 complex requires activation by specific N-blocked dipeptides (e.g., Z-Leu-Leu) to exhibit proteolytic activity *in vitro,* the mechanism of *in vivo* activation remains unclear.

In this study, we identified novel activators that enabled the structural determination of the ClpC1P1P2 complex, providing insights into its assembly. Furthermore, we discovered that trehalose - a key metabolite and molecular crowding agent in *Mtb*, significantly enhances the activity of both ClpC1P1P2 and ClpXP1P2 complexes without the need for activating peptides. Analytical ultracentrifugation revealed that trehalose promotes the formation of these active complexes, mimicking intracellular conditions. These findings propose a new model of Clp system activation *in vivo*, offering promising avenues for therapeutic targeting in tuberculosis treatment.

**Significance Statement:** The proteolytic complex formed by the essential proteins ClpP1 and ClpP2, along with their specific ATP-dependent activators ClpX and ClpC1, has emerged as a highly attractive target for anti-tuberculosis drug development. While previous studies have shown that ClpP1P2 can be activated *in vitro* by small peptide activators, its *in vivo* activation mechanism remains unclear. In this study, we identify novel activators and demonstrate that trehalose, a key metabolite in *Mycobacterium tuberculosis*, enhances ClpC1P1P2 and ClpXP1P2 activity without the need for activating peptides. These findings propose a new model for Clp system activation in *Mycobacterium tuberculosis*, advancing our understanding of its regulation and potential as a therapeutic target.

## Introduction

Tuberculosis, a leading cause of death from infectious diseases, claims nearly 2 million lives annually and affects one-third of the global population as a latent infection(1). Current treatments for tuberculosis are suboptimal, requiring prolonged regimens with multiple high-dose antibiotics, often leading to severe side effects and contributing to rising drug resistance (2). Moreover, *Mycobacterium tuberculosis* (*Mtb*) becomes increasingly resistant to available antibiotics. This underscores an urgent need for novel therapeutic strategies that can selectively target essential biological pathways in *Mtb*. We have characterized an enzyme representing such a target - the ClpP1P2 protease complex (3, 4). ClpP1 and ClpP2, both essential for *Mtb* viability and infectivity (3, 4), form a protease complex distinct from its mammalian counterparts. While mammalian cells harbor structurally different ClpP proteases in the mitochondrial matrix, no homolog exists in the cytosol, making the mycobacterial ClpP1P2 complex an attractive therapeutic target. ClpP is a highly conserved family of multimeric serine proteases originally discovered in *E. coli* (*5, 6*). ClpP homologs are present in a wide range of bacteria, as well as in chloroplasts and mitochondria in eukaryotes. The active enzyme is composed of two heptameric rings with 14 proteolytic sites located within the central chamber (7). Most microorganisms have a single *clpP* gene, while *Mtb* has two *clpPs, clpP1 and clpP2* (*3, 4*). The ClpP protease can only hydrolyze short peptides, whereas its complex form, in association with the AAA+ ATPases ClpC1 and ClpX, degrades proteins (its physiological function). AAA+ ATPases ClpC1 and ClpX, both essential for *Mtb* viability, selectively bind protein substrates, unfold them and translocate them into the ClpP proteolytic chamber for degradation (3, 4).

*In vitro*, recombinant *Mtb* ClpP1 or ClpP2 form homo-tetradecamers, but proteolytic activity is only achieved through the assembly of a heteromeric ClpP1P2 complex in the presence of specific activators, such as N-blocked dipeptides or derivatives, i.e. Z-Leu-Leu(4, 8). Z-Leu-Leu stimulates ClpP1P2 complex formation at relatively high concentrations (5 mM). Therefore, we set out to obtain more potent activators that could be used at lower concentrations. Using Z-Leu-Leu as a scaffold, we synthesized 81 N-protected dipeptides and evaluated their potency in activating *Mtb* ClpP1P2. Some of these newly synthesized molecules showed up to 20 times higher specific activity than Z-Leu-Leu. Notably, in the presence of one activator, Bz-Leu-Leu, we were able to determine the first structure of the ClpC1P1P2 complex.

Finally, we also addressed the question of how active Clp complexes are formed *in vivo*. We found that mimicking intracellular conditions using crowding compounds (such as trehalose or PEGs) stimulated the formation of active ClpC1P1P2 and ClpXP1P2 complexes even without activator peptides.

## Results

### Effect of N-terminal group (X_3_ position) on ClpP1P2 activation

Building on our previous findings identifying Z-Leu-Leu (X_3_-X_2_-X_1_) as an activator of ClpP1P2(4), we employed it as a scaffold to synthesize 81 new peptides, systematically varying the N-terminal group at the X_3_ position (for a complete list of compounds, the activation potency and abbreviations, see Table 1). These compounds were evaluated for their ability to activate ClpP1P2 using a peptidase assay with acetyl-PKM-7-amido-4-methylcoumarin (Ac-PKM-amc) as the substrate (8, 9). Activation potency was expressed as a percentage relative to Z-Leu-Leub at 4 mM, which was defined as 100% activity.

**Table 1.**
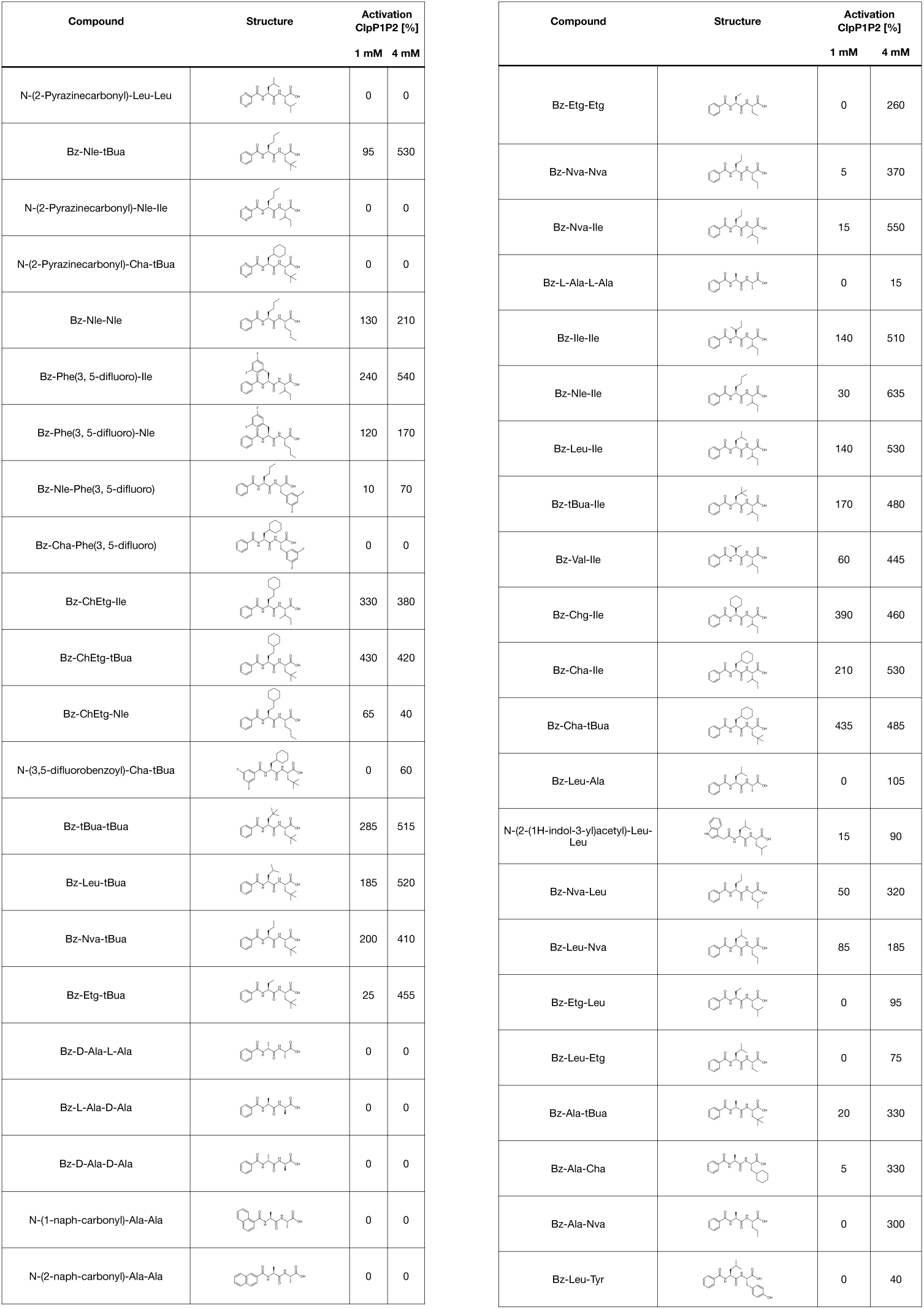
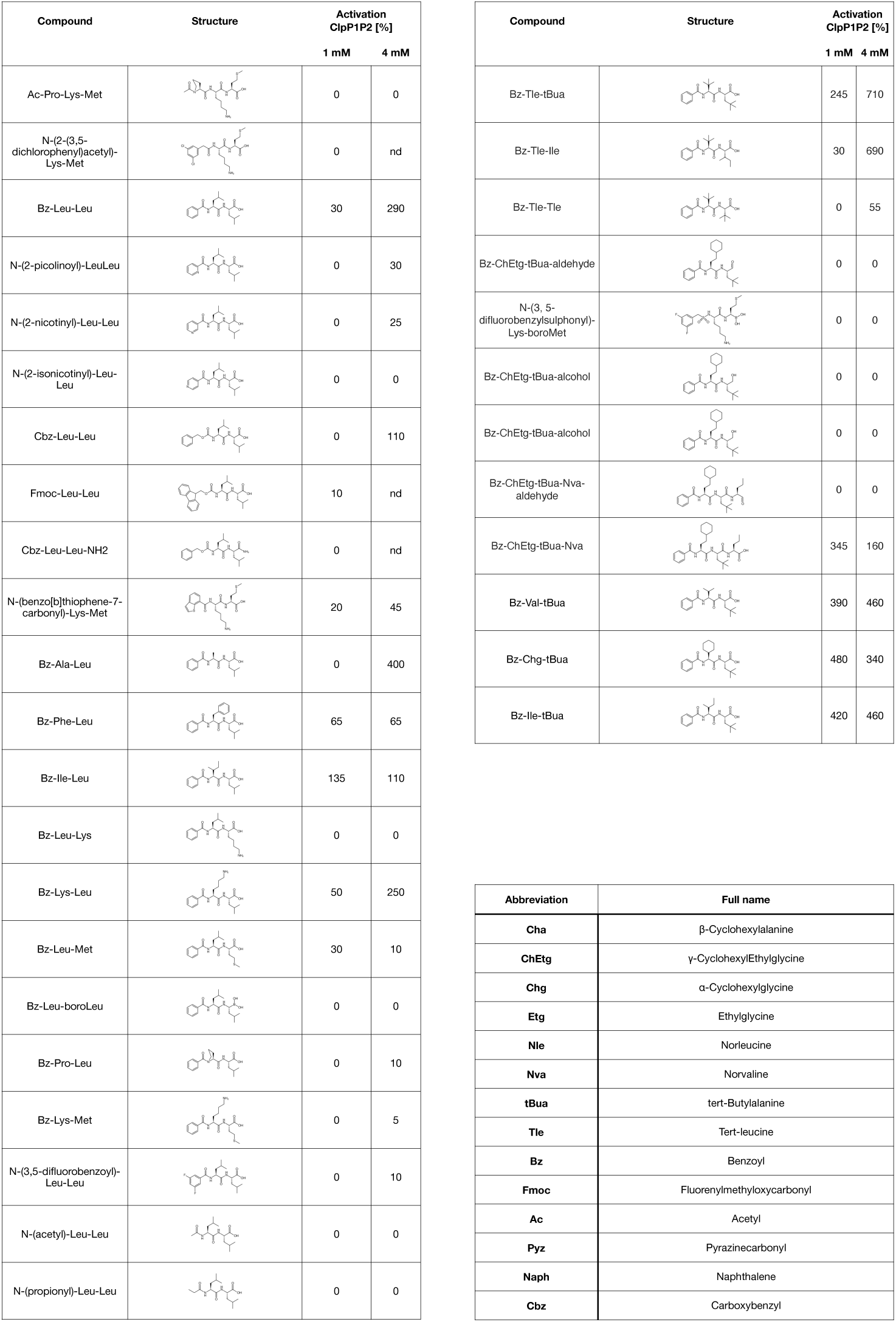
List of all peptide activators and their structure.

As a first step, we kept the peptide frame unchanged and tested how the substitution of the N-terminal carboxybenzyl group (Z) in X_3_ position with a variety of other N-blocked groups affected activation capacity (see Table 1). Most of the modified activators were either close in potency to Z-Leu-Leu (e.g. N-(2-1H-indol-3-yl-acetyl)) or markedly less active (Ac, Propyl, Isonicotinoyl, Fmoc or Pyz). The only blocking group that increased the potency of the scaffold was Bz (Fig 1A). As we reported previously, at 4 mM Bz-Leu-Leu was 3 times more active than Z-Leu-Leu(10). Interestingly, modifications of the Bz-group (i.e. addition of fluoride in N-(3,5 -difluorobenzoyl-Leu-Leu) markedly decreased the potency of the activator (Fig 1A). Based on these results, Bz-Leu-Leu was chosen as a new scaffold to determine the optimal residues for the X_2_ and X_1_ positions.

**Figure 1.**
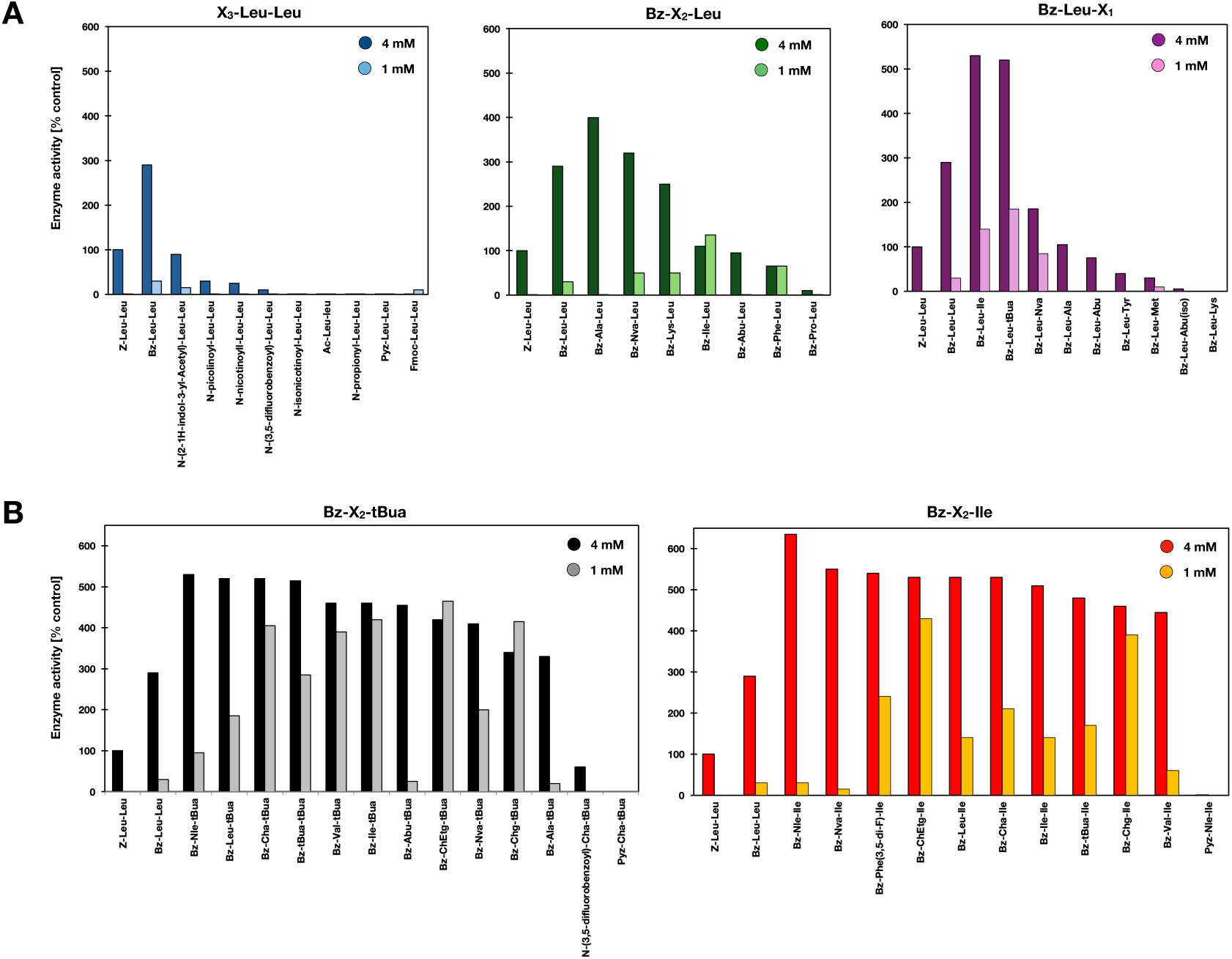
Action of novel peptide activators on *Mtb* ClpP1P2 and ClpP from other species. Action of different N-protected dipeptides on *Mtb* ClpP1P2 activity. Activation of *Mtb* ClpP1P2 was monitored continuously at a protein concentration of 40-100 nM using 20 μM Ac-Pro-Lys-Met-amc as a substrate. The compounds were tested at 1 and 4 mM concentration as described in Material and Methods. As comparison the activation level of the previously described and commonly used scaffold Z-Leu-Leu was taken as 100%. A) Modifications of the Z-Leu-Leu (X_3_-X_2_-X_1_) scaffold at X_3_ (left panel) and of the Bz-Leu-Leu (X_3_-X_2_-X_1_) scaffold at X_2_ and X_1_ positions (middle and right panel). B) Modification of the Bz-X_2_-tBua scaffold at X_2_ (left panel) and changes of Bz-X_2_-Ile scaffold at X_2_ (right panel).

### Determining optimal amino acids for X_1_ and X_2_ positions of Bz-X_2_-X_1_ activating peptide

Keeping the Bz in X_3_ and Leu in X_1_ positions, we searched for the best amino acids (or their derivatives) for the X_2_ position. As shown in Fig 1A, Ala was the best candidate for the X_2_ position, followed by Nva and Leu. At 4 mM, Bz-Ala-Leu activated ClpP1P2 better than Z-Leu-Leu or Bz-Leu-Leu.

Search for the optimal candidates for X_1_ position in Bz-Leu-[X_1_] identified Ile and tBua as best choices (Fig 1A). Bz-Leu-Ile and Bz-Leu-tBua at 4 mM stimulated ClpP1P2 peptidase activity 5-6 times better than Z-Leu-Leu at the same concentration (Fig 1A). Our results suggested that Ala in X_2_ and tBua/Ile in X_1_ position would constitute the best activator for ClpP1P2. Indeed, all compounds with tBua at X_1_ position were better activators than Z-Leu-Leu or Bz-Leu-Leu (Fig 1B). Compounds with tBua in X_1_ and either Cha, Chg, Val or Ile at the X_2_ position activated ClpP1P2 similarly at 1 mM and 4 mM. Bz-ChEtg-tBua and Bz-Chg-tBua showed higher activity at 1 mM than at 4 mM, indicating possible inhibition at high concentrations of activator (Fig 1B). Similar results were obtained for compounds with Ile at X_1_ position (i.e. Bz-ChEtg-Ile and Bz-Chg-Ile (Fig 1B) and no inhibition of peptidase activity was observed at high concentrations of these compounds (Fig 1B). In agreement with our prior observations, the substitution of Bz in X_3_ position with N-(3,5-difluorobenzoyl) or Pyz led to a loss of activity.

### New activators have higher potency and affinity to ClpP1P2 than Z-Leu-Leu

For all activators, which were significantly (8-27 times) more potent than Z-Leu-Leu, concentration dependence, Hill coefficients and K_D_s were determined. We have previously established that Z-Leu-Leu binds to ClpP1P2 in a highly cooperative manner with a K_D_ of 2.2 mM and Hill coefficient 5-6 (4). We found that several new activators (i.e. Bz-Leu-Leu, Bz-Nva-Ile, Bz-ChEtg-tBua, Bz-ChEtg-Ile, Bz-Cha-tBua and Bz-Ile-tBua) had higher affinity for ClpP1P2, with K_D_s 2-4 times lower than that of Z-Leu-Leu (Fig S1), while only small changes were observed for the number of molecules bound to the protease complex , with Hill coefficients in the range 4-6 (Fig S1).

### Structure of ClpC1P1P2 reveals an asymmetric assembly

As referred above, *in vivo* ClpP1P2 is not a peptidase, but rather a protease that catalyzes the degradation of protein substrates, which are recognized, unfolded (if necessary) and translocated into its chamber by ATPases like ClpC1 and ClpX. Therefore, to fully understand how this system works, structural information on the full active proteolytic complex in association with its cognate ATPases is of paramount importance. Previous studies have only characterized the individual parts of the complex and no structure exists of the full ClpC1P1P2 complex (10–12). While in the absence of activator ATPase-ClpP1P2 complexes are labile, we believed that using the peptide activators reported here could stabilize the complex sufficiently for cryo-electron microscopy (cryo-EM) structure determination. In addition, we introduced a stabilizing point mutation (F444A) in ClpC1 to favor its active hexameric state (12). To obtain an active complex, cryo-EM samples were prepared in the presence of ATPγS, peptide activator (Bz-Leu-Leu) and substrate (GFP-ssrA or FITC-casein). We decided to use Bz-Leu-Leu as it was one of the first new activators to be characterized and it has been successfully used by us to obtain the structure of the ClpP1P2 complex.

Both protein substrates, GFP and casein, yielded similar structural assemblies, with ClpC1 engaging the ClpP2 ring of the ClpP1P2 barrel (Fig 2A). Cryo-EM analysis revealed significant conformational heterogeneity, consistent with the dynamic nature of this molecular machine. As the majority of the substrate and the highly flexible NTD domains of ClpC1 were invisible to cryo-EM, we decided to only solve the structure of the GFP-ssra bound complex due to better data quality. As expected from a highly flexible molecular machine like the ClpC1P1P2 complex, the complex does not exist in a single static conformation. Indeed, it adopts multiple different conformations both, ClpC1 with respect to ClpP1P2 and within ClpC1 itself. This behavior is illustrated in supplementary Fig S2A, which shows 4 representative conformations, and Video 1, which shows a traverse trajectory across the latent conformational landscape encountered in the ClpC1P1P2 sample using DynaMight (13). While the ClpP1P2 structure seems unchanged in all cryo-EM maps, ClpC1 adopts several conformations on top of ClpP1P2, with the C-termini of ClpC1 and ClpP2 (shown in the inlet of Fig S2A) being either further apart or closer to each other. Similar conformational heterogeneity has been reported for the ClpC1P1P2 complex of *S. hawaiiensis* as well as other AAA+ unfoldases (14). Attempts to obtain a better insight into the engagement of the C-termini of ClpC1 and ClpP2 by masking failed, suggesting that this interaction is rather flexible or of a transient nature. In order to obtain the best structure of the ClpC1P1P2 complex for model building, we only solved the most abundant conformation, which was determined at a resolution of 3.09 Å. In order to improve resolution we employed masking of both the ClpC1 complex and the ClpP1P2 complex in a step-wise manner. The complete workflow can be seen in Table 2. The final high-resolution composite map (EMD-52840) was created by aligning the two focused maps of ClpC1 (EMD-52764) and ClpP1P2 (EMD-52766) against the unmasked consensus map of the ClpC1P1P2 complex (EMD-52765). The resulting structure (PDB 9IF4) shows an asymmetric hexameric ClpC1 on top of ClpP1P2 with a 26 residue peptide within the ClpC1 substrate pore, similar to other AAA+ ATPase-ClpP complexes (see Fig 2A). The asymmetric shape of ClpC1 is determined by the arrangement of the individual subunits and their pore loops in a typical staircase manner around the substrate, with one subunit (P6) completely detached from the substrate (Fig 2B/C). The ClpC1 subunits closer to the substrate (P2, P3, P4 and P5) are well resolved, while the detached subunits P6 and P1 have poor resolution, with the D2 domain only partially visible. The interaction of ClpC1 and ClpP2 is mediated by the LGF loops of ClpC1, with the Phe residue buried in the interface between two different ClpP2 subunits. Most of the LGF loop of ClpC1 is invisible, presumably due to high flexibility, apart from the LGF motif, which is likely tightly bound and well resolved for all 6 ClpC1 subunits. Due to the asymmetric fit of ClpP2 and ClpC1, one of the LGF loop binding sites on ClpP2 remains empty (see Fig 2D). All nucleotide sites in ClpC1 are occupied: the substrate-bound subunits (P2, P3, P4, P5) are bound to ATPɣS (both in the D1 and the D2 domain) and the detached subunits P1 and P6 to ADP (Fig 2E/F). Interestingly, while the X-ray structure of ClpP1P2 (PDB 5DZK) with Bz-Leu-Leu shows an even distribution of Bz-Leu-Leu in both ClpP1 and ClpP2 (10), our complete complex structure reveals asymmetric binding of Bz-Leu-Leu with preferential binding to ClpP2 (Fig 2G). Although residual density for Bz-Leu-Leu can be detected in ClpP1, the main binding site is in ClpP2, where all active sites are occupied with Bz-Leu-Leu and well resolved (Fig 2H). The orientation of Bz-Leu-Leu within ClpP2 is as described previously, with the benzoyl ring facing the S1 pocket and the catalytic residues Ser110 and His135 (10). The ClpP1P2 complex shows high similarity (RMSD 0.58 Å) to the structure previously solved by X-ray crystallography (10). The most striking differences are the N-termini of ClpP2, which form highly stable extended beta-hairpins in the X-ray structure but are invisible in our complex structure, probably due to high flexibility, likely reflecting the association with the highly mobile ATPase ClpC1 in our structure.

**Figure 2.**
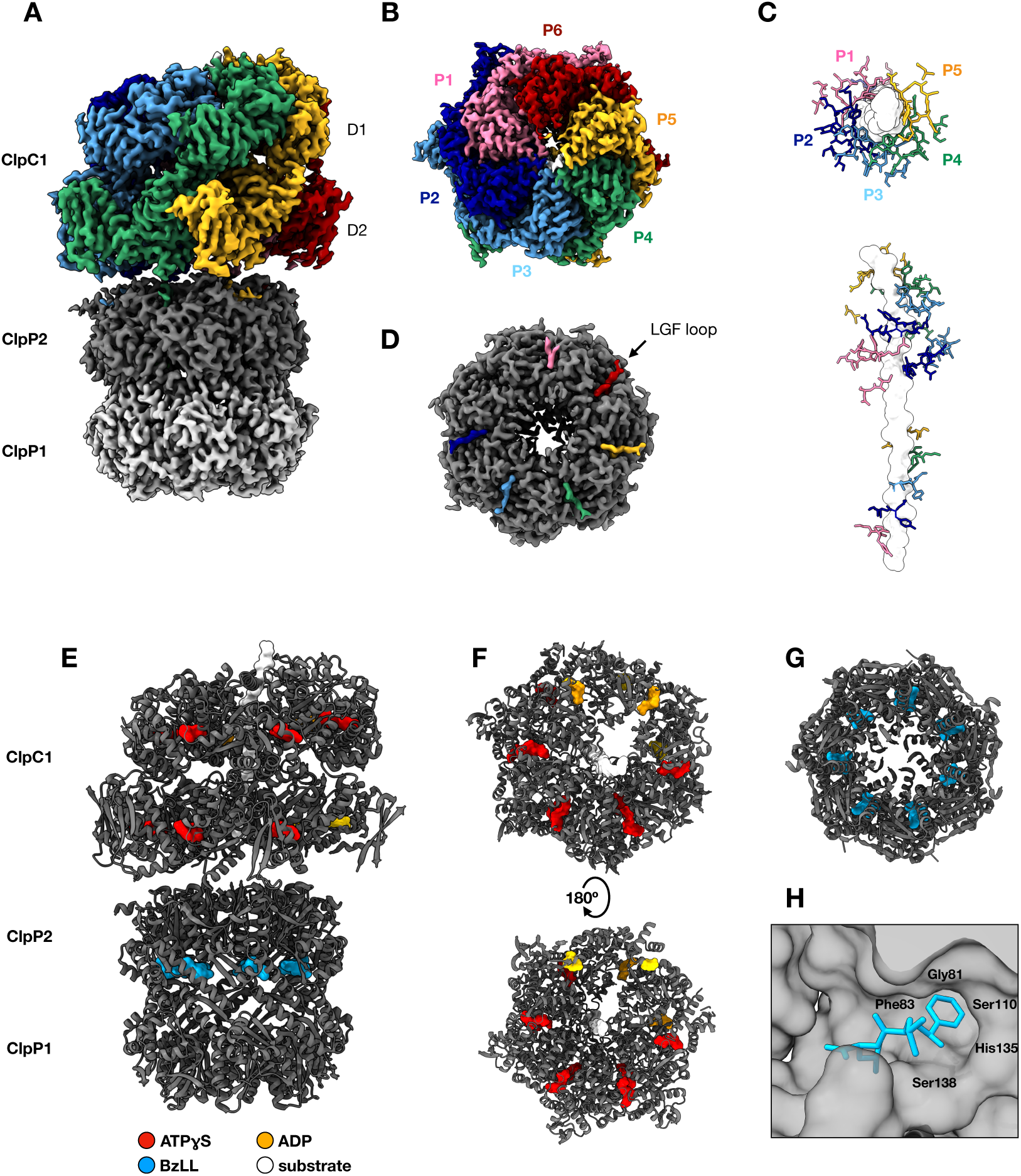
Structure of the *Mycobacterium tuberculosis* ClpC1P1P2 complex. A) Overall structure of the ClpC1P1P2 complex (PDB: 9IF4). The ClpC1 hexamer is depicted in rainbow, the ClpP2 heptamer in dark grey and ClpP1 in light grey. ClpC1 exclusively interacts with the ClpP2 heptamer. The interaction interface appears flexible with low resolution in the C-terminal loops of ClpP2 and the LGF loops of ClpC1, whereby only the LGF motif itself can be resolved within the binding pocket of ClpP2 (D). B) The ClpC1 hexamer shows the typical arrangement of a AAA+ ATPase where substrate bound protomers (P2-P5) show higher stability and substrate detached protomers higher flexibility (P1, P6). The interaction of the different pore loops of subunits P1-P5 can be seen in (C), while the pore loops of P6 are detached. D) The LGF motif in ClpC1 mediates the interaction with ClpP2. Due to the asymmetry in the number of subunits, one of the binding pockets in ClpP2 remains empty. E) Ligand binding in the ClpC1P1P2 complex. ATPγS is depicted in red, ADP in orange and Bz-Leu-Leu in cyan. F) All nucleotide sites in the ClpC1 D1 and D2 subunit are in the nucleotide bound state. Whereby the substrate bound ClpC1 protomers P2-P5 are bound to ATPγS, in both the D1 and the D2 subunit, and the detached protomers P1 and P6 are bound to ADP. G) Contrary to the previously published *Mtb* ClpP1P2 X-ray structure (PDB: 5DZK) Bz-Leu-Leu preferentially binds to ClpP2, with only residual binding to ClpP1. The orientation of Bz-Leu-Leu inside the binding pocket is as reported, with the Benzyl group orientated towards Ser110 (H).

**Table 2.**
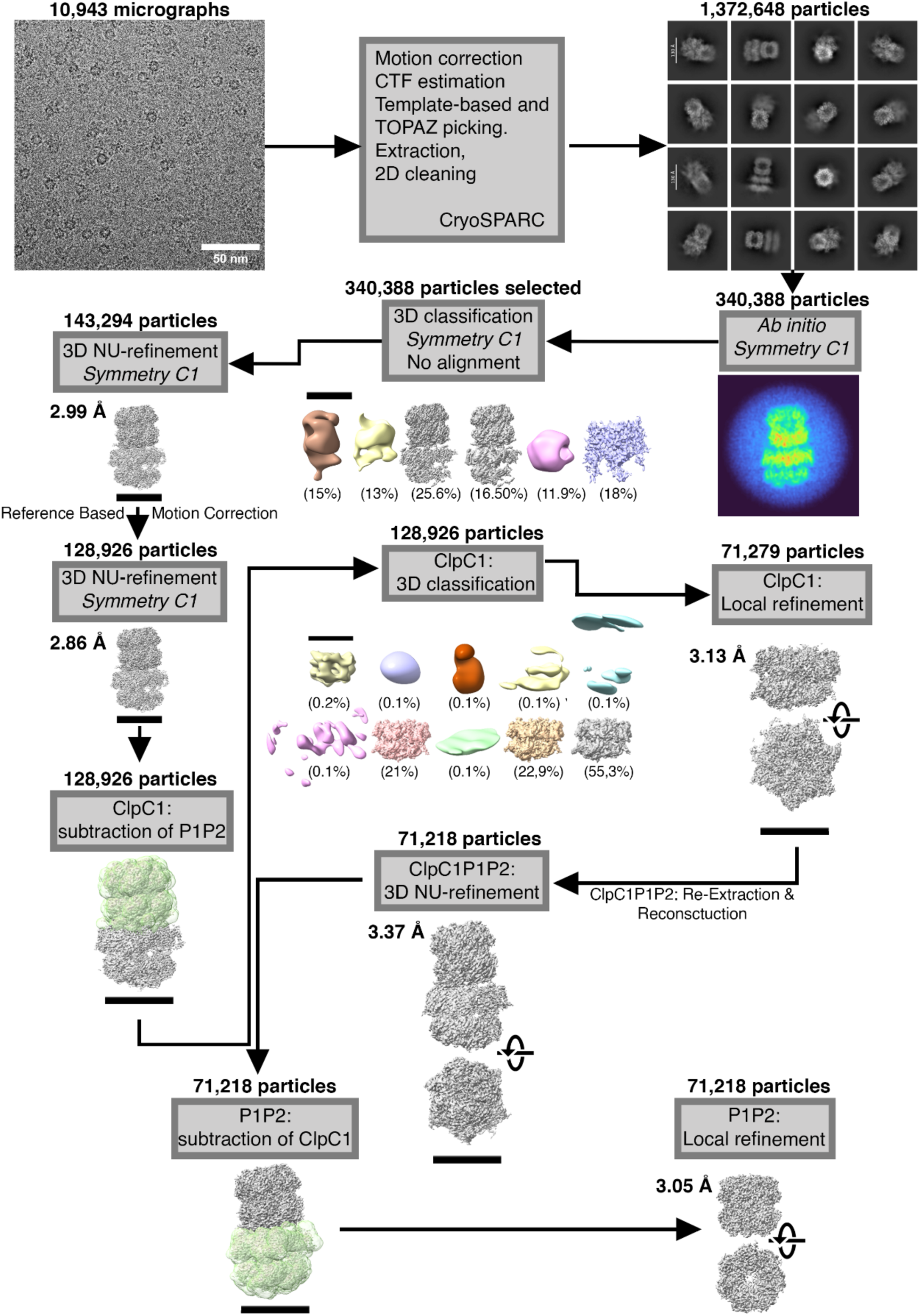
Pipeline for the cryo-EM structure determination of ClpC1P1P2.

Additionally, in our structure the usually unresolved C-termini of ClpP2 form alpha-helices pointing away from the ClpP1P2 barrel and are possibly in contact with the C-terminus of ClpC1 (see inlet of Fig S2A). Although the first 14 C-terminal residues of ClpP1 cannot be built into the cryo-EM map due to low resolution, the exit pore of ClpP1 appears to be closed in all our cryo-EM maps (Fig S2B). This contrasts the entry pore of ClpP2 that is always open (Fig S2C).

### Dipeptide activators act on human mitochondrial ClpP but not on homologous enzymes from other bacteria

We have previously demonstrated that dipeptide activators promote the association of ClpP1 and ClpP2 7-mers into active mixed ClpP1P2 complexes (4). Here, we tested whether these activators may have a similar effect on other ClpP complexes since all of them are built from two heptameric rings stuck together. Human mitochondrial ClpP (hClpP) mainly exists *in vitro* as inactive 7-mers and forms an active 14-mer only in the presence of the corresponding ATPase (15, 16). We investigated whether dipeptide activators can promote hClpP 14-mer formation in the absence of the ATPase. As shown in Fig 3A, in the presence of Bz-ChEtg-Ile or Bz-ChEtg-Bua activators hClpP peptidase activity increased more than 15 times, strongly suggesting that activators promote the association of inactive heptamers into tetradecamers. No such activation was observed when bacterial ClpPs from various sources (*S. aureus, E. coli* and *B. subtilis)* were tested (Fig 3B). On the contrary, some small inhibition of peptidase activity was seen for *E. coli* and *B. subtilis* ClpP. This seems logical since, unlike *Mtb* ClpP1P2 and hClpP, those bacterial enzymes exist in vitro as stable 14-meric structures and do not respond to the presence of activators. A similar result has been described previously when *Thermus thermophilus* ClpP was studied(17).

**Figure 3.**
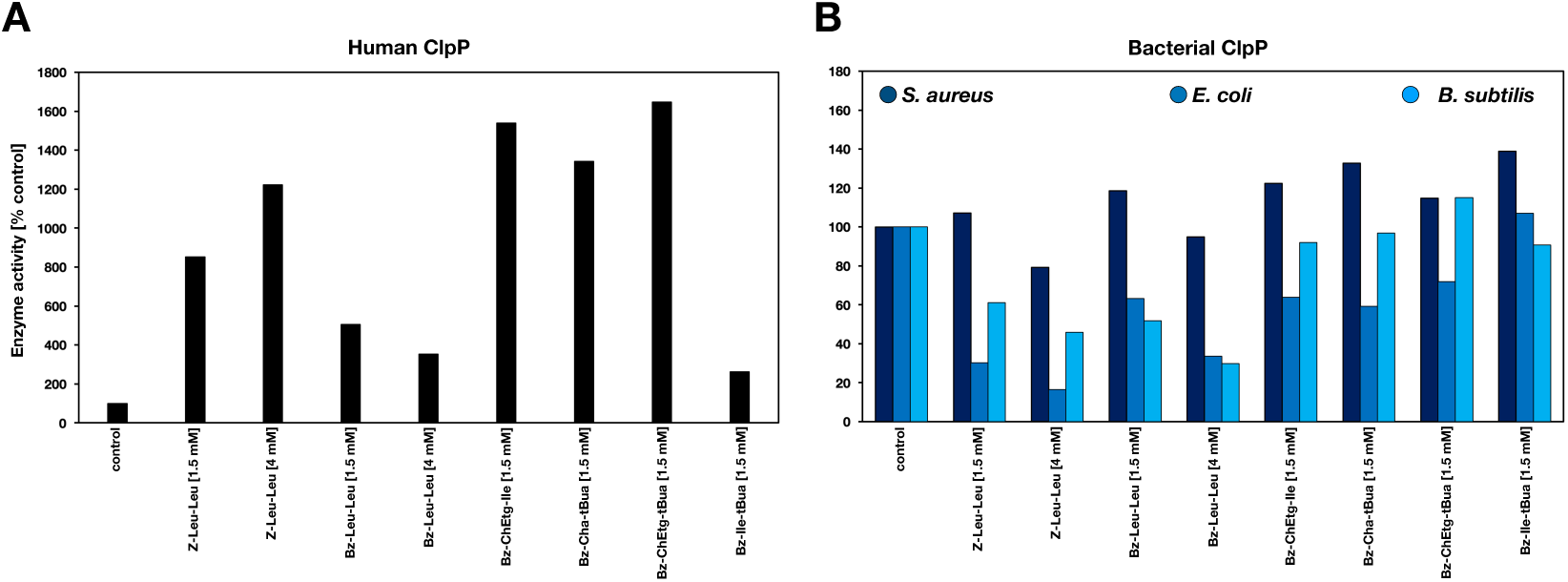
Action of novel dipeptide activator on ClpP from different species. The most potent activators were selected to test their effect on purified A) human ClpP and B) ClpP from different bacteria (*Staphylococcus aureus, Escherichia coli, Bacillus subtilis*). Z-Leu-Leu and Bz-Leu-Leu were used at 1.5 mM and 4 mM, all other activators were used at 1.5 mM. ClpP enzyme concentrations used were 180 nM (*S. aureus* and *B. subtilis*), 90 nM (*E. Coli)* and 200 nM (human).

### Dipeptide activators stimulate endogenous ClpP1P2 activity in cell extracts

We set to test whether dipeptide activators crucial for ClpP1P2 formation *in vitro* may also activate endogenous ClpP1P2. Cell extracts were prepared from *Mycobacterium smegmatis*, a non-infectious species closely related to *Mtb*, using a French press and cleared by centrifugation for 1 hour at 100,000 g. Basal ClpP1P2 peptidase activity was detected in the extract using Ac-PKM-amc as described (8). When some of the newly identified potent activators (Bz-Leu-Leu, Bz-ChgEtg-Ile, Bz-ChEtg-tBua and BzCha-tBua) were added to the extracts at 1.5 mM, endogenous ClpP1P2 activity increased up to 6-fold (Fig S3A). In agreement with our *in vitro* data (see above), new activators with the Bz-blocking group were 3 times more potent than Z-Leu-Leu (Fig S3A).

When either ClpP1 or ClpP2 was added to the cell extract together with activators, Ac-PKM-amc hydrolysis increased 2-3 fold over the levels observed with the activator alone (Fig S3B). These results indicate the formation of mixed ClpP1P2 complexes between the endogenous and exogenous *Mtb* ClpP1 and ClpP2, which is expected due to high homology between the species (91% for ClpP1 and 95% for ClpP2). Interestingly, adding ClpP1 or ClpP2 without activator did not increase endogenous ClpP1P2 activity.

To confirm that detected ClpP1P2-like activity is associated with high molecular weight complexes, the cell extract was fractionated using size exclusion chromatography on a Superdex 200 column and peptidase activity against Ac-PKM-amc was measured throughout the elution profile. Ac-PKM-amc hydrolyzing activity was associated only with high molecular weight fractions containing ClpP1P2, whose presence was confirmed by Western blot (Fig S3C, fractions 42 to 47). Notably, peptidase activity in these fractions increased upon addition of Bz-ChEtg-tBua (Fig S3D). These observations suggest that the detected endogenous peptidase activity could be attributed to *M. smegmatis* ClpP1P2 and that dipeptide activators can activate it. It was unclear however how ClpP1P2 complexes are formed *in vivo*, presumably, their formation is catalyzed by certain endogenous activators, but adding an exogenous synthetic activator further stimulates ClpP1P2 formation and/or activity.

### No low molecular weight activators of ClpP1P2 are found in *M. smegmatis* or *M. tuberculosis* extracts

While we found a way to activate ClpP1P2 *in vitro*, it is unclear whether similar mechanisms exist *in vivo*. We investigated whether any low molecular weight activators similar to compounds active *in vitro* (peptides or other metabolites) are present in cell extracts. The low molecular weight (LMW) fraction was obtained by filtration of concentrated *M. smegmatis* extracts. When the LMW fraction was added to the ClpP1P2 peptidase activity assay mixture, no increase in enzymatic activity was observed. These results suggest that *M. smegmatis* cytosol does not contain LMW activators for ClpP1P2 activity. We also tested LMW weight fractions from several other species of Mycobacteria *(M. tuberculosis, M. avium, M. abscessus, M. fortuitum, M. kansasii and M. chelonae)* obtained by aqueous or ethanol/acetic acid extractions and kindly provided by Dr. Clardy, HMS. None of the tested fractions could stimulate the activity of recombinant ClpP1P2 *in vitro* or endogenous ClpP1P2 in *M. smegmatis* extracts. On the contrary, some of the extracts partially inhibited ClpP1P2 activity. These findings indicate that ClpP1P2 activation *in vivo* may occur via mechanisms distinct from those observed *in vitro*.

### Trehalose, a natural molecular crowding agent, supports the activity of ClpXP1P2 and ClpC1P1P2 complexes in the absence of dipeptide activator

Since we did not find evidence of low molecular weight activators *in vivo*, we sought to mimic the intracellular environment rich in osmolytes and macromolecular crowders. Many organisms accumulate osmolytes in response to environmental stress, which serve as molecular crowders within the cytosol. As our team and others demonstrated, microorganisms typically accumulate the non-reducing disaccharide trehalose (α-d-glucopyranosyl-1,1-α-d-glucopyranoside) in such conditions (Kandror et al. 2002, 2004). In mycobacteria, trehalose is best known for its role as a structural component in cell wall glycolipids, including trehalose dimycolate (the cord factor), which has significant structural and immunomodulatory functions. Recent studies using metabolomics and an *in vitro* biofilm model of *Mtb* persistence revealed that *Mtb* persisters remodel trehalose metabolism to drive both transient drug tolerance and permanent drug resistance - an adaptive response referred to as the trehalose catalytic shift (Lee et al. 2019). Furthermore, research has shown that macromolecular crowding, particularly in environments with high concentrations of sugar-based solvents like trehalose, sucrose, sorbitol, and glycerol, enhances the catalytic efficiency of certain enzymes *in vitro*, with trehalose being the most effective sugar. Additionally, macromolecular crowding can promote protein-protein interactions, leading to oligomer formation (Mittal et al. 2015). Given these findings, and considering that ClpP1P2 and its associated unfoldases are exposed to high trehalose concentrations *in vivo*, we decided to investigate whether trehalose could enhance the enzymatic activity of Clp complexes *in vitro* in the absence of dipeptide activators.

Enzymatic activities of ClpP1P2, ClpX, ClpC1, ClpC1P1P2 and ClpXP1P2 were tested in the absence or presence of trehalose. No peptidase activity could be detected for ClpP1P2 in the absence of dipeptide activator whether trehalose was present or not (Fig 4A). Similarly, trehalose did not affect ATPase activity of ClpX (Fig 4A) or ClpC1 (not shown). Interestingly, peptidase activities of the ClpXP1P2 and ClpC1P1P2 complexes were markedly enhanced in the presence of trehalose in a concentration-dependent manner (Fig 4A,B,C). The effect of trehalose became evident at 0.05 M and reached maximum at 0.4-0.8 M. The maximal peptidase activity in the presence of trehalose is about 1/3 of the activity with dipeptide activator Bz-Leu-Leu (Fig 4D). It is noteworthy that trehalose also activated endogenous ClpP1P2 peptidase activity in *M. smegmatis* extracts when added at concentration of 0.8 M (Fig S4A). We also tested several other disaccharides instead of trehalose; while some had similar effects, trehalose was the most effective (not shown).

**Figure 4.**
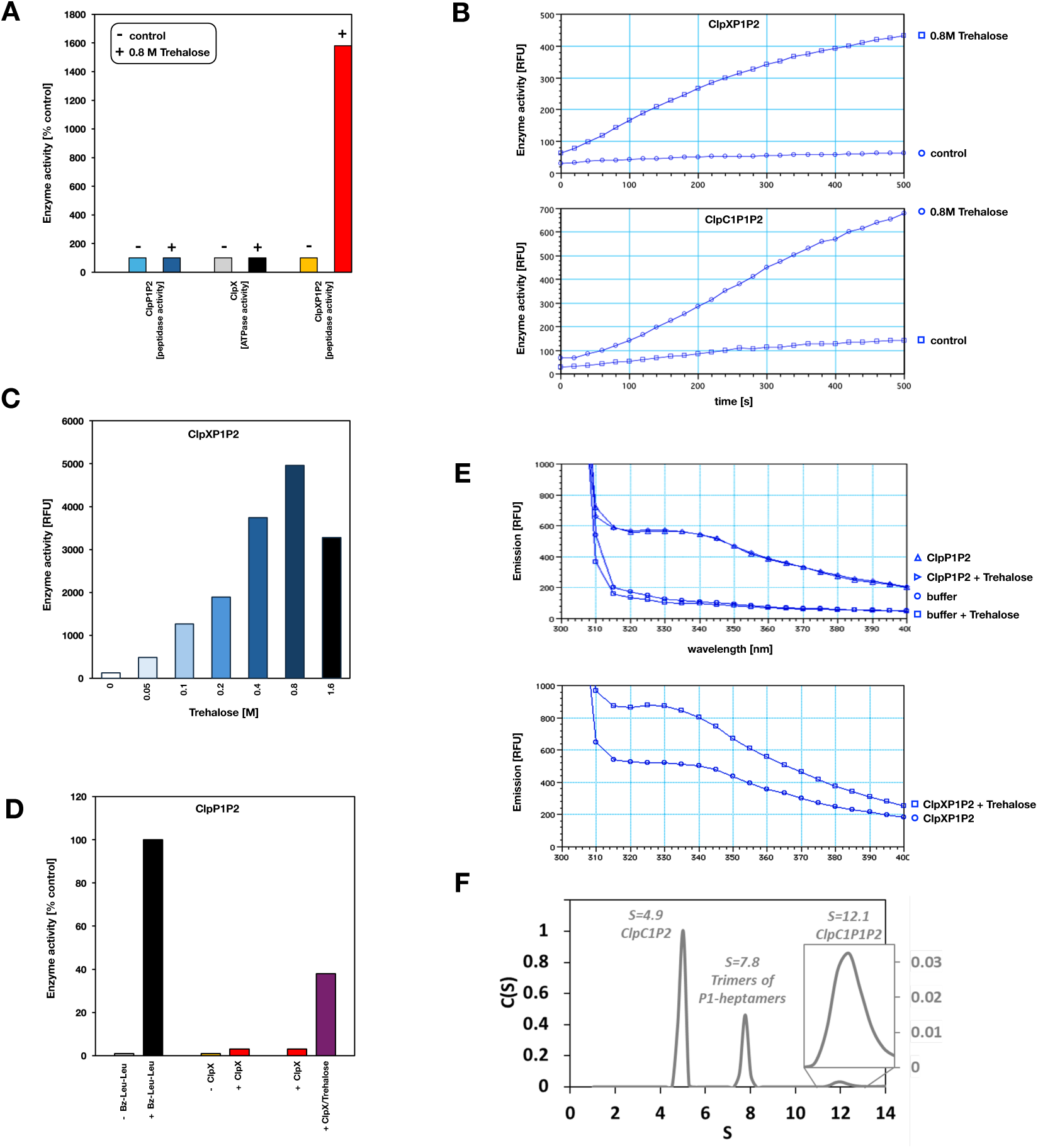
Effect of Trehalose on ClpP1P2 activity. A) Effect of trehalose (0.8 M) on ClpP1P2 peptidase, ClpX ATPase and ClpXP1P2 peptidase-activities. Activities in the absence of trehalose were taken as 100 %. B) Activation of ClpXP1P2 (top) and ClpC1P1P2 (bottom) peptidase activity with 0.8 M trehalose. C) Trehalose increases ClpXP1P2 peptidase activity with a maximum activation at 0.8 M trehalose. D) Peptidase activity of ClpP1P2 in the presence of Bz-Leu-Leu, ClpX and ClpX plus trehalose (0.8 M). Enzyme activity in the presence of Bz-Leu-Leu was taken as 100 %. E) Emission spectra of ClpP1P2 with ClpX in the absence and presence of trehalose. F) ClpC1P1P2 complex formation in trehalose solution analyzed by analytical ultracentrifugation (AUC). Truncated sedimentation distribution highlighting the most abundant species of the ClpC1-F444A, ClpP1 and ClpP2 mixture. An insert shows the peak corresponding to the ClpC1P1P2 complex. Experimentally derived sedimentation coefficient values - Svedberg units [S] - are shown above each peak. The full analysis can be found in Figure S5.

**Figure 5.**
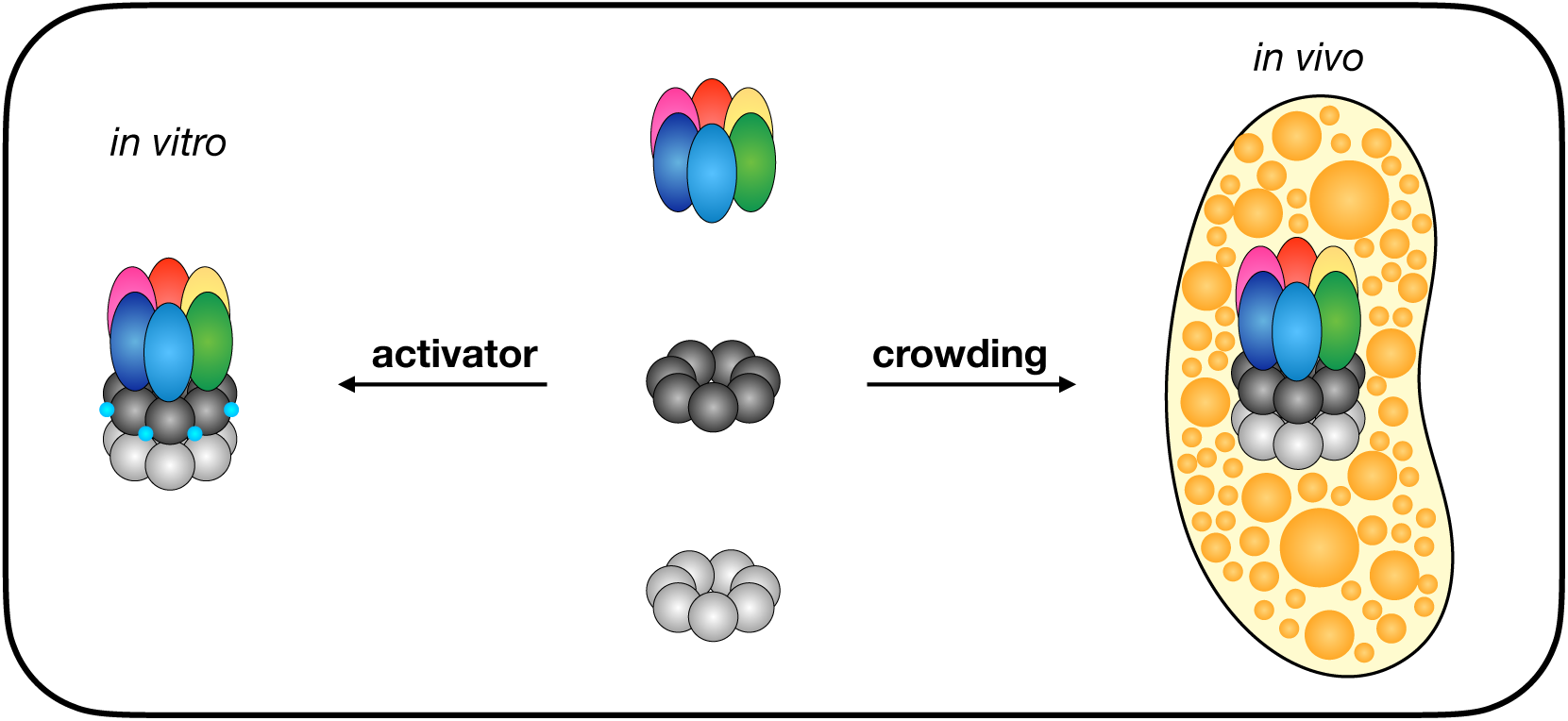
Formation of the active ClpC1P1P2 complex *in vivo* and *in vitro*. Formation of the active ClpP1P2 hetero-tetradecamer *in vitro* can be induced by N-blocked dipeptides such as Bz-Leu-Leu. No such low-molecular weight activators could be found *in vivo*. Here we show that complex formation through molecular crowding could be the main mechanism of activation in the cell. ClpC1 is shown in rainbow colors, ClpP2 in dark grey, ClpP1 in light grey and the activator in cyan.

To confirm that the trehalose effect on ClpXP1P2 and ClpC1P1P2 activity was due to its crowding effect, we used synthetic high molecular weight polymers - polyethylene glycols (PEG) of varied lengths (400-20,000 Da). PEG solutions of the same percentage (w/v) were prepared to keep the number of ethylene glycol groups the same. As shown in Fig S4B, the peptidase activity of ClpXP1P2 increased up to 10-fold in the presence of PEG, regardless of the length of the PEG chain (at the same w/v concentration). Similar results were obtained for ClpC1P1P2. At the same time, no change in the peptidase activity of ClpP1P2 or the ATPase activity of ClpX or ClpC1 was observed in the presence of PEG. Crowding agents seem to stimulate interaction between ClpP1P2 peptidase and ClpX and ClpC1 ATPases or stabilize this large complex, preventing its dissociation.

### Trehalose favors formation of large ClpXP1P2 and ClpC1P1P2 complexes

It has been reported that ClpP1P2 complexes can form *in vitro* in the absence of a dipeptide activator but much less efficiently than in the presence of an activator (18). In our hands, in the absence of a dipeptide activator, ClpP1P2 activity is negligible but increases significantly upon addition of the corresponding ATPases ClpX and ClpC1, still reaching only a fraction of peptidase activity obtained in the presence of the dipeptide activators (Fig 4D). In our experience, large ClpXP1P2 and ClpC1P1P2 complexes are transient and unstable, even in the presence of dipeptide activators, thus the crowding effect of trehalose could be beneficiary for their stabilization (4). To test this hypothesis, we attempted to test directly whether more complexes could be detected in the presence of trehalose. We used an assay based on changes in the fluorescence spectrum of Trp residues in the ClpP1P2 complex in response to conformational changes caused by the binding of ATPases (4). ClpP2 and ClpX do not have any Trp residues, and the only Trp in ClpP1 is present in residue 172 (Fig S4C). As we have previously shown, this residue is in the vicinity of the key aspartate in the protease catalytic triad. It is therefore a perfect probe for conformational changes, such as the ones promoted with activator binding. ClpC1 cannot be used for this assay as it contains multiple tryptophanes.

As shown in Fig 4E, trehalose by itself did not cause any changes in the Trp fluorescence of ClpP1P2. On the contrary, when ClpX was also present, a significant increase in Trp fluorescence was observed. Thus, while trehalose does not affect ClpP1 and P2 interaction, it favors the formation of ClpXP1P2 complexes. These data suggest that in a crowded intracellular environment, the formation of active ClpXP1P2 or ClpC1P1P2 proteolytic complexes may occur without any additional small molecule activators.

In order to demonstrate the formation of ClpC1P1P2 in the presence of trehalose, it was fundamental to resolve the complex from the remaining species, namely free ClpP1, ClpP2, ClpC1 and different partial complexes. One approach is to use size exclusion chromatography (SEC) but the use of high concentrations of trehalose leads to high pressure within the column, making it unsuitable for this study. Moreover, SEC has multiples disadvantages, including sample dilution, assumptions about the molecules shape and stationary phase interaction. Analytical ultra centrifugation (AUC) is a uniquely useful tool to investigate macromolecular characteristics such as size, shape, and stoichiometry. Unlike other biophysical approaches, where signals from all the species are superimposed, ultracentrifugation combined with a dual UV/visible and interference detection system enables hydrodynamic separation of species in solution according to their sedimentation coefficient (S). Thus, macromolecular properties can be probed without the need for standards, assumptions, or physical matrices, reflecting the kinetic and equilibrium properties of the interacting system in complex solution as close to a native environment as possible. Particularly in this study, AUC allowed us to observe the behavior of isolated or interacting systems in buffer containing trehalose. As shown in Fig 4F, the effect of trehalose on a mixture of ClpC1, ClpP1 and ClpP2, monitored by AUC, results in the formation of 3 main peaks at 4.9 S, 7.8 S, and 12.1 S, corresponding respectively to ClpC1P2, trimers of P1-heptamers and ClpC1P1P2 complex. This complex is formed of six ClpC1, seven ClpP1, and seven ClpP2 molecules, as expected. The sedimentation coefficient of the ClpC1P1P2 complex is close to the value calculated using hydropro (19) (12.4 S) based on the cryo-EM structure. It is likely that the formation of the complex induced by trehalose is a stepwise process with the prior formation of an intermediate composed of ClpC1 and P2. The full AUC profile of the complex and the individual proteins can be found in Fig S5.

## Discussion

The use of experimental models to study and explain biological phenomena is fundamental to modern science. Biochemical knowledge, in particular, has been significantly advanced through data obtained from *in vitro* models, including purified enzymes and cofactors in dilute solutions. While these models offer reliable, reproducible data, caution is necessary when extrapolating *in vitro* findings to biological systems. Although *in vitro* conditions are valuable for studying the structural and functional organization of protein machines, results from such studies must account for the fact that these systems operate within cells in a complex, crowded environment. These microenvironments can influence the activity and localization of proteins and other macromolecules involved in essential processes, thereby acting as non-specific modulators of bacterial cellular functions.

One example of a supramolecular machine influenced by these factors is the ClpC1P1P2 system in *Mtb*. The ClpP system in *Mtb* exhibits unique characteristics, starting with the involvement of two genes in forming the protease. Although other species, such as *Listeria monocytogenes* and *Streptomyces hawaiiensis*, also possess two different types of subunits, the heterogeneous ClpP1P2 complex in *Mtb* does not spontaneously form active complexes *in vitro* (4). Active complexes were first described when it was discovered that small peptides - initially expected to act as inhibitors - could instead activate the complex (4). These activators bind to the active site and induce allosteric activation across remaining active sites, explaining the observed activation (17, 20). Similar conformational changes have been observed in the ClpP of *Thermus thermophilus*, and we here suggest that active site activators might induce analogous changes in human ClpP (17). Additionally, we report a new advancement in developing more potent activators, which were tested and selected. These molecules are valuable tool compounds for biochemical research and potential drug development. Indeed, by utilizing Bz-Leu-Leu, we present the first cryo-EM structure of the *Mtb* ClpC1P1P2 complex. Consistent with previous genetic and biochemical studies, only the ClpP2 interface binds to ClpC1, forming an asymmetric complex with unique properties. It remains unclear why ClpP2 is preferred or the functional or physiological relevance of this feature, though a similar asymmetric interaction has been observed in the ClpCP1P2 structure of *Streptomyces hawaiiensis*. In our density maps, weak interactions between ClpP2’s C-terminal and ClpC1 in certain conformations may suggest that this interface is more stabilized. However, our model’s low resolution precludes a more definitive statement. Other structural factors may also explain the asymmetric arrangement. A related study on the *Mycobacterium smegmatis* ClpP1P2 complex found that regulation might also involve a ClpP1 C-terminal extension that blocks the chaperone binding pocket in ClpP1. Despite the genetic proximity, this C-terminal extension is absent in *Mtb* ClpP1.

Another intriguing question raised by the asymmetric structure is its functional significance. In the ClpC1P1P2 complex, ClpP1 adopts an extended conformation with an active site geometry. However, its apical pore is obstructed by N-terminal residues, making the release of products inside the cavity unclear. Additionally, the activator binds preferentially to ClpP2, with minimal binding observed in ClpP1’s heptamer. However, our data and previous studies indicate that ClpP1P2 complex formation functions as a regulatory switch for the complex. This may be especially relevant in *Mtb*, where the abundance of intrinsically disordered proteins would make them easy targets for an active protease.

While activators facilitate structural studies *in vitro*, they do not entirely explain how complex activation occurs *in vivo*. One possibility is that small, endogenous activators facilitate activation within cells, though we have not observed any activation by cell extracts from *M. smegmatis* or other mycobacteria. It remains possible that such activators are either unstable or produced only under specific cellular conditions. Alternatively, other mechanisms may be at play. In vitro assays typically use dilute protein solutions, but the cellular milieu is complex, crowded, and rich in sugars and osmolytes. Trehalose, a major component in *Mycobacterium*, can account for circa 70% of low molecular weight components in acid-dormant *M. smegmatis* (21). Notably, when trehalose is added to dilute solutions of ClpP1 and ClpP2, larger molecular weight species are promoted. This is consistent with molecular crowding promoting larger assemblies and among the species identified using AUC, about 3% corresponded to the ClpC1P1P2 complex (see insert Fig 4F).

Interestingly, the ClpC1P2 complex was also observed, suggesting it might act as a precursor to the ClpC1P1P2 complex, where ClpP2 binds ClpC1 prior to ClpP1 binding (Fig 4F). The activating effect of trehalose does not result from active ClpP1P2 complex formation, as it does with peptide activators, but rather depends on the interaction between ClpP1P2 and ClpC1. Importantly, the activation observed in the presence of trehalose is significantly lower than that achieved with the new active site activators, indicating two independent ClpP activation mechanisms: one mediated by trehalose promoting unfoldase assembly to ClpP1P2 and another through activators binding directly to ClpP1P2 active sites and inducing allosteric activation.

A final question to address is the *in vivo* relevance of trehalose activation. Assuming a 1.1 density for a water-trehalose solution, the 0.6 M trehalose used in our AUC assays corresponds to approximately 19% (w/w), but the exact intracellular trehalose concentrations in *Mtb* are difficult to determine and likely vary based on cellular conditions. Nonetheless, given the cytoplasm’s complexity, perhaps this exercise is futile, as no *in vitro* conditions can fully replicate cellular conditions. The cytosol behaves as a nonideal solution, where thermodynamic activity depends not merely on individual concentrations but on factors such as high concentrations of diverse macromolecules, reduced free water availability, and complex spatial organization. Our data suggest that molecular crowding, whether promoted by trehalose or other high molecular weight molecules, e.g., proteins and DNA, facilitates ClpC1P2 and ClpC1P1P2 complex formation. It seems very reasonable to assume that cytoplasmic crowding would exceed that induced by a 0.6 M trehalose solution, supporting the likelihood of this mechanism *in vivo*.

In summary, *in vitro* studies have significantly advanced our understanding of the ClpC1P1P2 complex in *Mycobacterium tuberculosis*, particularly the mechanisms underlying small-molecule activation. However, they fall short of fully capturing the complex’s *in vivo* activation. This study provides new insights into the natural assembly of the ClpC1P1P2 complex in the absence of exogenous activators. The observed asymmetric structure and regulatory mechanisms reflect specialized adaptations to *Mtb*’s crowded cytoplasmic environment, highlighting the importance of studying these systems under conditions that closely mimic physiological contexts. These findings have direct implications for drug development, especially given the limited success of past screening efforts targeting the *Mtb* ClpP system. Refining screening methodologies to incorporate conditions that simulate the intracellular environment could significantly enhance the discovery of effective therapeutic agents.

## Materials and Methods

### Synthesis of the *M. tuberculosis* ClpP1P2 protease activators

Synthesis of the N-acylated dipeptides and peptidomimetics N-acyl-X_2_-X_1_ were accomplished by acylation of the commercial or in-house made dipeptides and peptidomimetics X_2_-X_1_ with the corresponding acyl chloride in aqueous NaHCO_3_ solution or organic solvent (i.e. DCM) with DIEA as the base (Scheme 1). The target compounds were purified by recrystallization or by reverse-phase (RP)-HPLC using a Varian semi-preparative system with a Discovery C18 569226-U RP-HPLC column, and the mobile phase was typically made by mixing water (0.1% TFA) with acetonitrile (0.08% TFA) in gradient concentration. Purities determined by HPLC analysis were >95%.

**Scheme 1.**
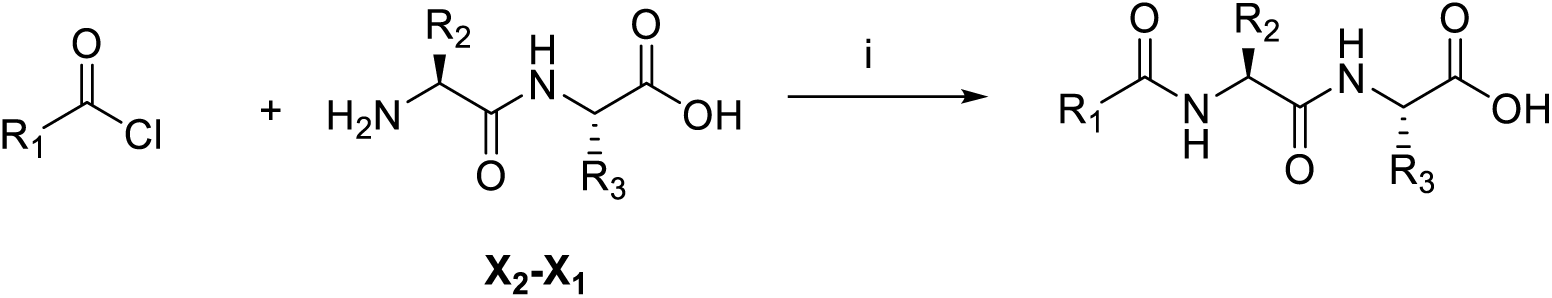
Reagents and conditions: i: aq. aqueous NaHCO_3_; or DIPEA, DCM.

<u>Exemplary experimental procedure for the preparation of *N*-Bz-Leu-Leu:</u>

To a suspension of commercially available L-Leu-L-Leu-OH (24.4 g, 100 mmol) in THF (450 mL) was added saturated aqueous NaHCO_3_ (320 mL). The mixture was cooled to 0°C, and benzoyl chloride (15 ml, 130 mmol) was added dropwise. The cool bath was removed, and the mixture was stirred at room temperature overnight and then acidified with 3N HCl to pH 3, extracted with ethyl acetate. The organic layer was washed with brine, condensed, and the residue was purified by recrystallization from EtOAc/n-Hexane. The obtained crystal was dissolved with 60% acetonitrile in water and lyophilized to give the product Bz-Leu-Leu (22 g, 63%) as a white powder. ^1^H NMR (D_2_O-CD3CN, 0.85:0.15) δ 0.90 -1.05 (m, 12H), 1.70 - 1.85 (m, 6H), 4.40 - 4.45 (m, 2H), 7.40 - 7.85 (m, 5H). MS (ESI^+^) m/z (rel intensity): 349.2 ( [M + H]^+^, 100), 218.1 ([M - Leu]^+^, 30).

### Bacterial strains, plasmids, expression, growth of cells

As described before, all *Mtb* proteins were expressed in the *E. coli* BL21 strain lacking endogenous ClpP and ClpX. All proteins had C-terminal 6His-tags except ClpX, which contained an N-terminal 6xHis-tag. A truncated form of ClpX (residues 60-426) was used throughout this study.

### Purification of *Mtb* enzymes

The purification of recombinant *Mtb* ClpP1 and ClpP2 was carried out as described previously [1]. The *Mtb* ClpX and ClpC1 were purified at 4° C using buffer B containing 50 mM Tris-HCl pH 7.6, 50 mM KCl, 0.1 mM DTT, 1 mM Mg-ATP and 10% glycerol. Cells were resuspended in two buffer volumes and lyzed with a French Press at 1500 psi. The extract was centrifuged at 100, 000 x g and mixed with Ni-NTA agarose. After 4 h incubation, Ni-NTA agarose resin was transferred to a column, and proteins were eluted using step gradient (25, 50, 100 and 200 mM) of imidazole in buffer B. The active fractions containing nearly homogeneous proteins were combined and concentrated to 1-3 mg/ml by Millipore MWCO 10 kDa cut filter and fractioned further by gel filtration on a column (2.5 x 22 cm) of Sephacryl S-300 equilibrated in the same buffer. The protein peak was collected, concentrated to ∼ 3 mg/ml and stored at – 80° C. All enzymes purified migrated as a single band in the SDS PAGE.

### Peptidase Assay

All assays of peptidase activity were performed at 37° C in black 384-well plates using a Plate Reader SpectraMax M5 (Molecular Devices, USA). 2-3 min were allowed for the reaction mixtures to reach 37°C. Each well contained 20 mM of fluorogenic peptide, 20-50 nM ClpP1P2 in 80 ml of buffer A (20 mM phosphate buffer pH 7.6 with 100 mM KCl, 5% glycerol and activators). DMSO concentration never exceeded 2 %. The reaction was initiated by adding the enzyme and peptidase activity, followed by monitoring the production rate of fluorescent 7-amino-4-methylcoumarin-amc from peptide-amc substrates at 460 nm (Ex at 380 nm). The deviation of fluorescence value in three independent measurements was not more than 5%.

### Activator Assay

Usually, to check the activation of enzymes, we mixed inactive ClpP1 and ClpP2 in the presence of activators and preincubated them at room temperature for 1h. Previous experiments indicate the max activation can be reached at about 30 min preincubation. The specific activation of newly synthesized compounds is calculated using the following formula: maximum activation (%)/concentration of compound used. The activity of Z-Leu-Leu used as a scaffold at 4 mM was taken as 100 %.

### Proteinase assay

*Mtb* ClpP1P2 was also assayed in 96-well plates using the fluorescent protein substrates GFPssrA and FITC-casein. To measure GFPssrA degradation by the ClpXP1P2 complex, each well contained 2 mM Mg-ATP, 500 nM GFPssrA, 75-100 nM ClpP1P2 and 300-400 nM ClpX hexamer in 100ml of buffer A. GFPssrA fluorescence was measured at 510 nm (Ex at 470 nm). To measure FITC-casein degradation by ClpC1P1P2, each well contained 2 mM Mg-ATP, 150-200 nM ClpP1P2, 500-700 nM ClpC1 hexamer and 1-1.2 mM FITC-casein in 100 ml of buffer A. FITC-casein fluorescence was measured at 518 nm (Ex at 492 nm). The deviation of fluorescence value in three independent measurements was not more than 10%.

### ATPase assay

ATP hydrolysis was measured with the enzyme-linked assay using pyruvate kinase and lactic dehydrogenase (PK/LDH). 2 µg of pure ClpC1 or ClpX was mixed with 100 µl of the assay buffer (50 mM Tris-HCl pH 7.8; 50 mM KCl; 10% glycerol; 1 mM phosphoenolpyruvate; 1 mM NADH; 2 units of pyruvate kinase/lactic dehydrogenase (Sigma); 4mM MgCl_2_ and 1 mM ATP) and the ATPase activity was followed by measuring the coupled oxidization of NADH to NAD spectrometrically at 340 nm. Measurements were performed in triplicate, which agreed within 5 %.

### Determination of Hill Coefficients

The Hill coefficients and K_D_s were calculated using the ReaderFit software (Hitachi Solutions America, Ltd. 2014). The equation used is the 4-parameter logistic (4PL) nonlinear regression model, which was used to fit data points:

(X)= ((A-D)/(12+((X/C)^B)))+D

Where X represents the used concentration and F(X) is the response value. A stands for the minimum asymptote, which is the responsive value. B is the slope factor and represents the Hill coefficient, indicating the cooperation between compound X and the Enzyme. C is the inflection point; at this point, the curve changes its direction and thereby equals the K_D_ of the underlying enzymatic reaction (half maximum activity at a given enzyme concentration). D is the maximum asymptote, which indicates the maximum enzyme activity at the given enzyme concentration in this case. Activation of *Mtb* ClpP1P2 was monitored continuously using as substrate 20 μM Ac-Pro-Lys-Met-amc and 50-100 nM ClpP1P2 in the presence of 1 and 4 mM compounds as described in Material and Methods. For comparison, the activation level previously described and used as scaffold rZ-Leu-Leu was taken as 100 %.

### Analytical Ultra-Centrifugation (AUC)

ClpC1-F444A (2.7 µM), ClpP1 (2.7 µM), ClpP2 (2.7 µM) and the subsequent complexes (made of 2.7 µM of each partner) diluted in HEPES 50 mM pH 7.4, NaCl 100 mM and MgCl_2_ 10 mM solution supplemented with Trehalose 0.6 M and ATPψS 1 mM were centrifuged at 42,000 rpm in an Optima AUC analytical ultracentrifuge (Beckman Coulter), at 20°C in an 8-hole AN 50–Ti rotor equipped with 12-mm double-sector aluminium epoxy centerpieces. Detection of the biomolecule concentration as a function of radial position and time was performed by interference detection. Sedimentation velocity data analysis was performed by continuous size distribution analysis c(s) using Sedfit 16.36 software (22). All the c(s) distributions were calculated with a fitted fractional ratio f/f0 and a maximum entropy regularization procedure with a confidence level 0.95. Buffer viscosity and density were measured on a Viscosizer TD and Anton Paar DMA 5000M respectively. Partial specific volumes were also determined with Sednterp.

### Cryo-EM sample preparation

Cryo-EM grids of ClpC1P1P2 (1 µM), Bz-Leu-Leu (1 mM), ATPγS (1 mM) and GFP-ssra or FITC-casein (10 µM) were vitrified using a Vitrobot Mark IV (Thermo Fisher Scientific). Quantifoil Cu/Rh 1.2/1.3 300 mesh grids were previously glow-discharged for 30 s at 25 mA. Aliquots of 3 *μ*l of the prepared sample were added onto the grids, blotted for 3 s at 4°C and 95% humidity, and plunged into liquid ethane.

### Cryo-EM data collection

Screening and data acquisition for the ClpC1P1P2 complex sample was performed using a 200 kV FEI Talos Arctica equipped with a Falcon 4i direct electron detector (Thermo Fisher Scientific). A total of 10942 movies were acquired at a nominal magnification of 120,000× (corresponding to a pixel size of 0.84 Å/pixel), with a defocus range of −0.8 to −2.2 *μ*m. Movies were fractionated to 666 frames with a total exposure of 2.17 s and a total electron dose per movie of 46 e^−^/Å^2^.

### Image analysis

The acquired cryo-EM dataset was processed using cryoSPARC v4.5.3+240807 (Patch). A total of 10,943 electron event representation (EER) movies, collected using Aberration-Free Image Shift (AFIS) and containing beam shift information, were imported and preprocessed. Motion correction was applied using a patch-based algorithm to correct beam-induced drift, followed by patch-based contrast transfer function (CTF) estimation for each micrograph in cryoSPARC (23) . Micrographs with a CTF fit resolution beyond 6 Å were excluded, resulting in 10,608 high-quality micrographs for further analysis.

Initial processing started by picking using crYOLO (24) and TOPAZ (25) inside Scipion (26) . Coordinates from both pickers were merged, removing duplicates and picks in the carbon foil with Deep Micrograph Cleaner (27) , adding a total number of 2,657,864 coordinates. These were extracted with 100 pixels box size at 3.12 Å/px, and 2D-classified with cryoSPARC for several rounds, resulting in 410,151 particles. Next, *ab initio* reconstruction and 3D heterogeneous refinement into 4 classes yielded a set of 340,120 particles. These were reextracted at 1.56 Å/px and box size 200 and subjected to Non-uniform refinement, followed by 3D classification into 10 classes, without recomputing alignments, and with solvent mask applied on ClpC1 (initialization mode: PCA). Particles from each output class were indepentently subjected to an additional round of Non-uniform refinement. Four of these maps were selected in Fig. S2 to illustrate conformational heterogeneity of the whole complex. Additionally, one of these classes (30,404 particles) was fed as input for DynaMight (13) to further explore its conformational variability, despite having been previously 3D-classified. For that, particles were first imported in Relion-5.0 (28) and auto-refined enabling Blush regularization. Next, particles were subjected to DynaMight, using 16,000 Gaussians. A set of volumes spanning a horizontal trajectory across the low-dimensional representation of the latent space was used for Supplementary Video 1.

A more thorough reprocessing pipeline was followed to obtain the high resolution maps in Fig. 2. Particle picking was conducted using a combination of template-based and TOPAZ machine-learning protocols to maximize particle identification accuracy. The resulting datasets were merged, duplicates were removed, and 1,372,648 particles were extracted using a box size of 600 pixels. To streamline processing and reduce computational demands, particles were Fourier-cropped to 200 pixels.

Multiple rounds of 2D classification were performed to refine the stack further, resulting in 340,388 particles. These particles were then used for ab-initio 3D reconstruction.

Subsequent 3D classification was conducted using six classes with the most complete ab-initio result. Two of these 3D classes, representing the most complete and well-defined assemblies, were combined and subjected to homogeneous refinement, followed by another round of ab-initio 3D reconstruction and classification. A final stack of 128,926 particles was utilized for Non-Uniform refinement, resulting in a consensus map with a resolution of 2.99 Å based on the FSC (Fourier Shell Correlation) curve at the 0.143 criterion and postprocessing with an automatically estimated *B*-factor of -81.1. To enhance 3D map quality, Reference Based Motion Correction was followed by another round of Non-Uniform refinement with the 3D map resolved at 2.86 Å based on the FSC curve at the 0.143 criterion and postprocessing with an automatically estimated *B*-factor of -80.8. To analyze each protein unit of the ClpC1P1P2 complex and to clarify the resolution of each one, ClpC1 and P1P2 were subtracted and refined further. The ClpC1 part of the final 3D map was subtracted and underwent Local refinement, resulting in a resolution of 3.15 Å based on the FSC curve at the 0.143 criterion and postprocessing with an automatically estimated *B*-factor of -95.2. The corresponding particles underwent 3D classification with ten classes. The best 3D class containing 71,279 particles proceeded further to Local refinement, resulting in a 3D map with a resolution of 3.13 Å (EMDB: EMD-52764) based on the FSC curve at the 0.143 criterion and postprocessing with an automatically estimated *B*-factor of -79.7. Those particles with the corresponding transformation matrix were re-extracted (71,218), and the ClpC1P1P2 3D volume was reconstructed.

One more cycle of Non-Uniform refinement was performed with the best 71,218 particles using the previously reconstructed 3D volume and its mask as a reference. The resulting 3D map of ClpC1P1P2 was obtained at a resolution of 3.37 Å (EMDB: EMD-52765) based on the FSC curve at the 0.143 criterion and postprocessing with an automatically estimated *B*-factor of -82.0. From this 3D map, the P1P2 part of the structure was subtracted and underwent Local refinement, resolving at a resolution of 3.05 Å (EMDB: EMD-52766) based on the FSC curve at the 0.143 criterion and postprocessing with an automatically estimated B-factor of -77.7.

No symmetry constraints were imposed during refinements, preserving the asymmetric features inherent to the ClpC1P1P2 complex.

### Model Building

Model building was performed using the software Coot. To obtain the highest possible resolution the two focused maps of ClpC1 (EMD-52764) and ClpP1P2 (EMD-52766) were used and aligned against the full complex map (consensus map) of ClpC1P1P2 (EMD-52765) using the software ChimeraX to create the final high-resolution composite map for model building (EMD-52840). Several refinement cycles were performed using the software package Phenix and the software package CCP4 for ligand building. The resulting structural model was deposited on the protein data base (PDB) with the accession number 9IF4.

## Acknowledgments

This work benefited from access to the Instruct-ERIC centers CNB-CSIC, Madrid, for cryo-EM sample preparation and data collection and the Instruct Image Processing Centre. Financial support was provided by Instruct-ERIC (PID 26164 and 30641). Additionally, this project has received funding from the European Union’s Horizon 2020 research and innovation programme under grant agreement No 101004806 providing access to the MOSBRI TNA site Institut Pasteur through the project MOSBRI-2023-183. This work was also supported by FCT - Fundação para a Ciência e a Tecnologia, I.P., through MOSTMICRO-ITQB R&D Unit (DOI 10.54499/UIDB/04612/2020; DOI10.54499/UIDP/04612/2020) and LS4FUTURE Associated Laboratory (DOI10.54499/LA/P/0087/2020).

This work is dedicated to the memory of Prof Alfred Goldberg (1942–2023), a pioneer in the study of protein degradation, in particular ClpP protease, and most of all a unique mentor and scientist.

He was involved at the genesis of this research, and we are very sad he is not with us to see it published.

## Figures and Tables

**Figure S1.**
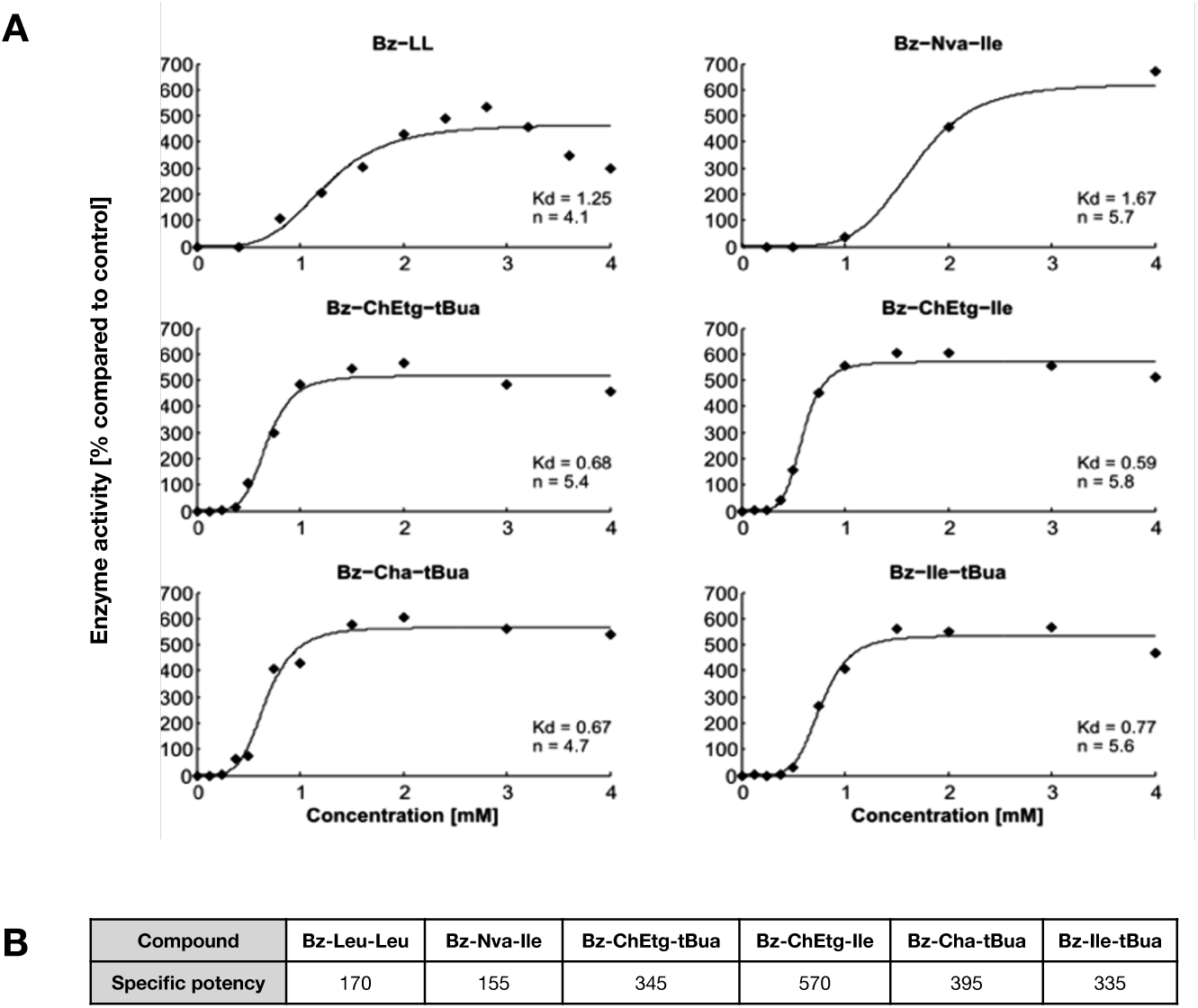
Determination of Hill-coefficients and K_D_ (mM) for six selected compounds. A) The following activators: Bz-Leu-Leu, Bz-Nva-Ile, Bz-ChEtg-tBua, Bz-ChEtg-Ile, Bz-Cha-tBua and Bz-Ile-tBua were chosen to determine K_D_ and Hill-coefficients (n number) using 100 nM of ClpP1P2 and 20 μM Ac-Pro-Lys-Met-amc. Calculations were done with the Reader Fit software (HitachiSolutions America LTD 2014) as described in Materials and Methods. B) Specific potency of activators calculated as ratio: % of maximum activation / concentration of compound. The Z-leu-Leu activator, used as a scaffold, has a specific potency equal to 18.

**Figure S2.**
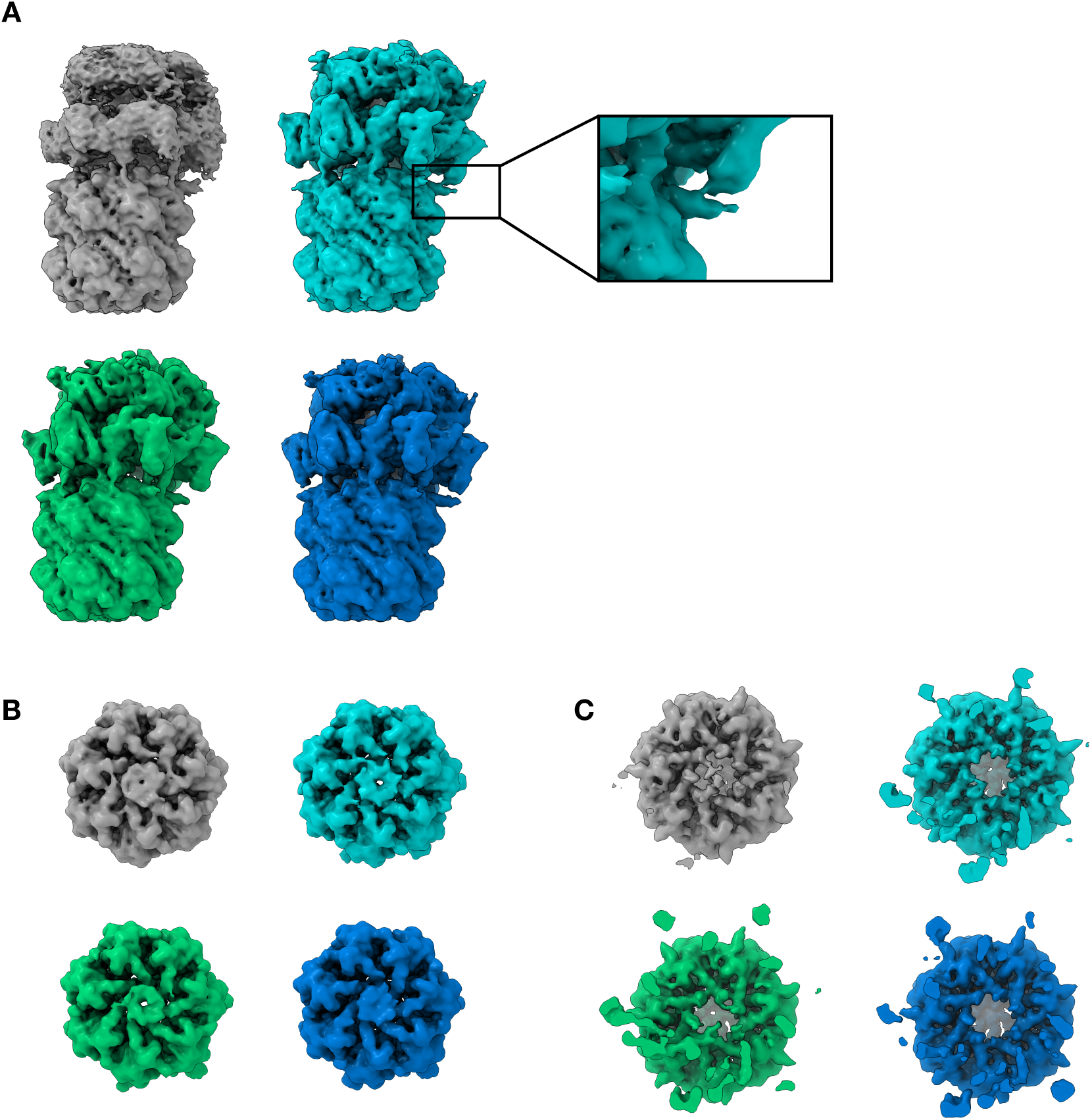
Conformations of the ClpC1P1P2 complex observed by cryo-EM Multiple conformations were observed in the ClpC1P1P2 cryo-EM sample. Here we show 4 representative conformations (A). A main difference is the proximity of the C-termini of ClpP2 and ClpC1 (see inlet in cyan) in the 4 maps. The same behaviour can be observed in Video1. Comparison of the presumable exit (B, ClpP1) and entry (C, ClpP2) pores of the ClpP1P2 barrel within the ClpC1P1P2 complex formed by the respective N-termini.

**Figure S3.**
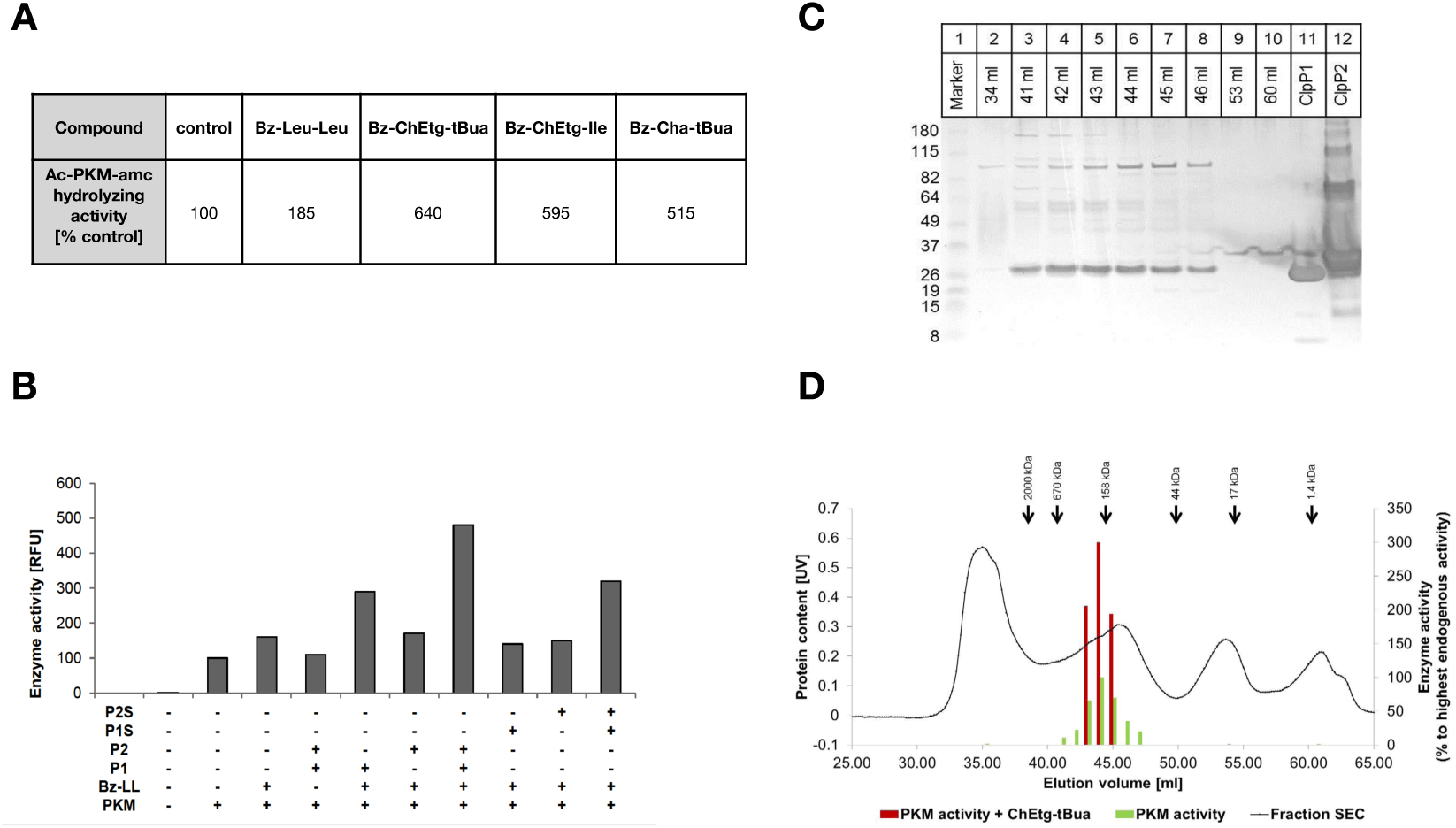
Endogenous ClpP1P2 activity of a *Mycobacterium smegmatis* extract. A) Action of different dipeptide activators on endogenous ClpP1P2 from *M. smegmatis* cell extracts. B) 10 μl of *M. smegmatis* extract were used to examine the endogenous ClpP1P2 activity in the presence of 140 nM purified enzyme (ClpP1, ClpP2 and their catalytically inactive mutants ClpP1S and ClpP2S) and 2.5 mM Bz-Leu-Leu activator. Activity of 10 μl *M. smegmatis* extract in the absence of additives (enzymes or activator) was taken as 100%. D) 500 μl of *M. smegmatis* extract were run on a Superdex 200 10/300 column. Elution volumes of molecular weight markers are indicated by an arrow: Vitamin B12 (1.4 kDa), myoglobin (horse, 17 kDa), ovalbumin (chicken, 44 kDa) γ-globulin (bovine, 158 kDa), thyroglobulin (bovine, 670 kDa) and blue dextran (2000 kDa). Fractions were collected in 500 μl steps. Of each fraction 20 μl were taken to assay enzyme activity with 20 μM Ac-PKM-amc. The same fractions were used for Western Blot analysis using anti-His antibody (C).

**Figure S4.**
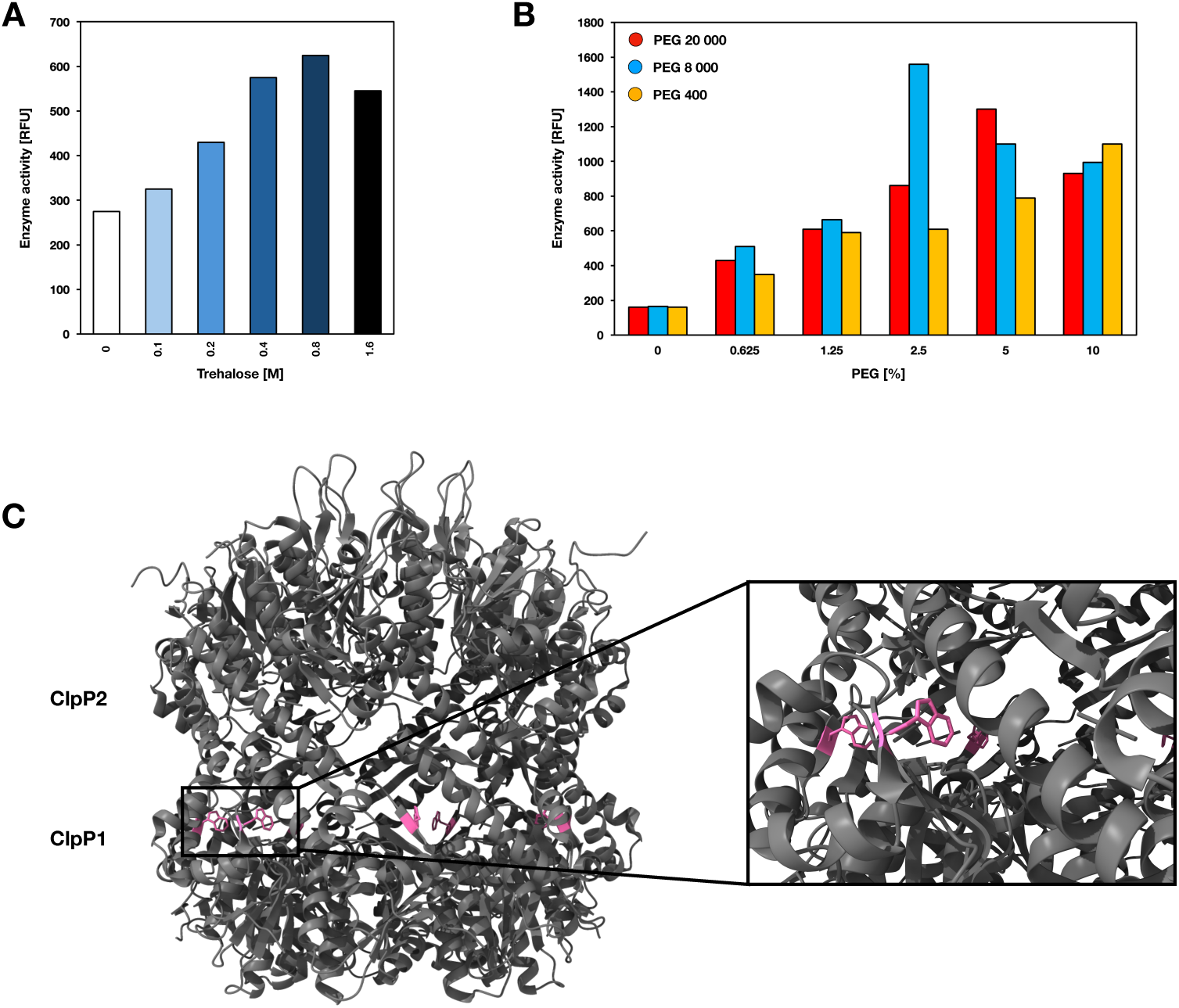
Molecular crowding as an activator of ClpP1P2 activity A) Trehalose increases Ac-PKM-amc hydrolyzing activity of *M. smegmatis* extracts with an activation maximum at 0.8 M trehalose. B) ClpXP1P2 peptidase activity in the presence of PEGs with different molecular weights (red: 20 000, blue: 8 000, yellow: 400) and concentration (0-10 %). C) Localization of the single tryptophane residue in ClpP1 (PDB: 5DZK, Trp colored in pink). Both ClpX and ClpP2 possess no tryptophane residues.

**Figure S5.**
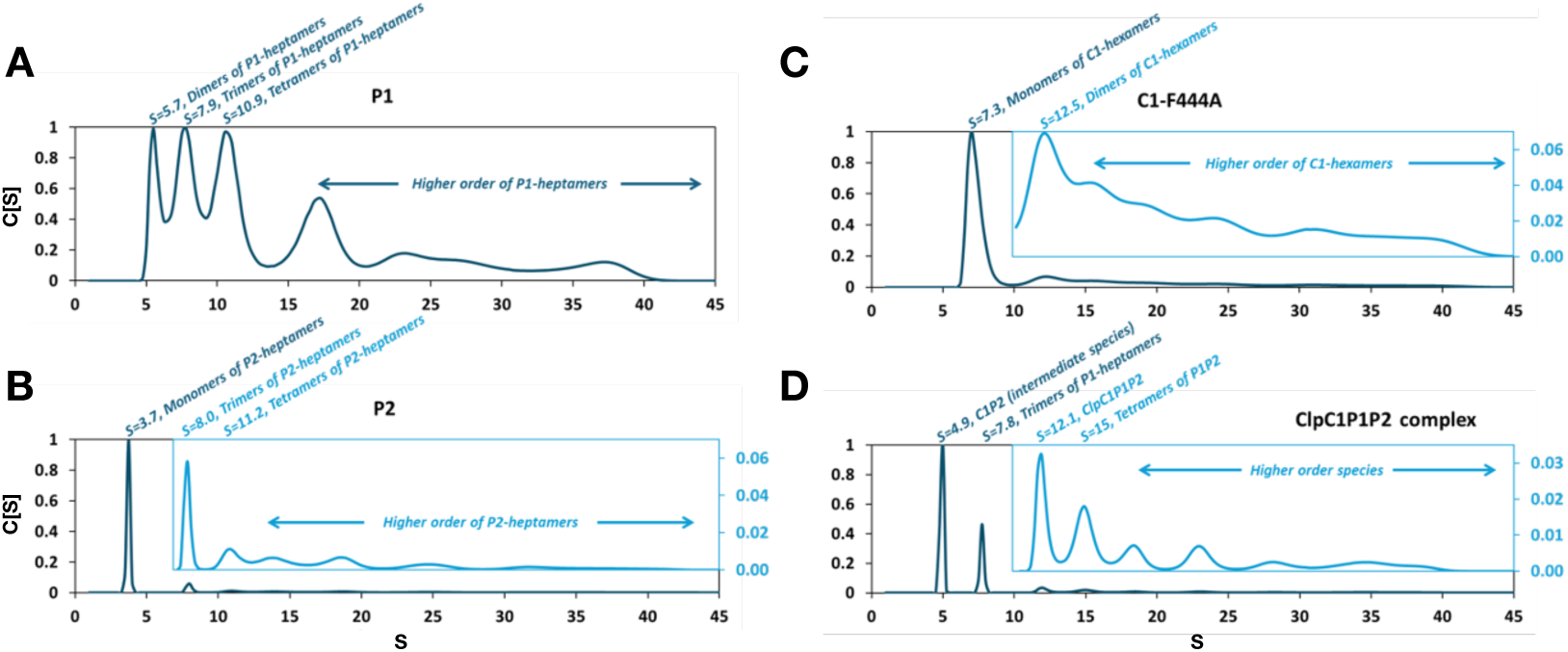
Effect of Trehalose on oligomerization and the formation of the active ClpC1P1P2 complex. Analysis of the ClpC1P1P2 complex formation in trehalose solution by analytical ultracentrifugation (AUC). Measurements were carried out on ClpP1 (A), ClpP2 (B), ClpC1-F444A (C) and ClpC1P1P2 (D) complex. All low-concentration oligomers and intermediate species are zoomed with a light blue insert. Experimentally derived sedimentation coefficient values (Svedberg units [S]) are shown above each peak and have a 0.2 standard deviation. Theoretical sedimentation coefficients were calculated from the crystal structure (5DZK PDB) using Hydropro 10 (García De La Torre et al) with a hydrated radius of 3.1 Å for the atomic elements. The theoretical values of the monomers of ClpP1-heptamers, ClpP2-heptamers and ClpC1-hexamers as well as the ClpC1P1P2 complex were calculated to be 3.9, 3.9, 9.8 and 12.6, respectively. The measured and theoretical values of the ClpP1-heptamers and ClpP2-heptamers monomers are in good agreement. In the case of ClpC1-hexamers and the ClpC1P1P2 complex, it should be noted that 1800 amino acids of each ClpC1-hexamer are missing from the PDB structure, which explains the differences between measured and calculated values.

**Video 1. Structural diversity of the ClpC1P1P2 complex**

